# PyEvoCell: An LLM- Augmented Single Cell Trajectory Analysis Dashboard

**DOI:** 10.1101/2024.11.21.624686

**Authors:** Sachin Mathur, Mathieu Beauvais, Arnau Giribet, Nicolas Aragon Barrero, Chaorui-Tom Zhang, Towsif Rahman, Seqian Wang, Jeremy Huang, Nima Nouri, Andre Kurlovs, Ziv Bar-Joseph, Peyman Passban

## Abstract

**Motivation:** Several methods have been developed for trajectory inference in single cell studies. However, identifying relevant lineages among several celltypes is a challenging task and requires deep understanding of various celltype transitions and progression patterns. Therefore, there is a need for methods that can aid researchers in the analysis and interpretation of such trajectories.

**Results:** We developed PyEvoCell, a dashboard for trajectory interpretation and analysis that is augmented by large language model (LLM) capabilities. PyEvoCell applies the LLM to the outputs of trajectory inference methods such as Monocle3, to suggest biologically relevant lineages. Once a lineage is defined, users can conduct differential expression and functional analyses which are also interpreted by the LLM. Finally, any hypothesis or claim derived from the analysis can be validated using the veracity filter, a feature enabled by the LLM, to confirm or reject claims by providing relevant PubMed citations.

**Software Availability and Implementation:** The software is available at https://github.com/Sanofi-Public/PyEvoCell. It contains installation instructions, user manual, demo datasets, as well as license conditions (including limitation to non-commercial uses only).

**Supplementary information:** Supplementary information is attached.

## Introduction

Trajectory inference (TI) analysis is crucial to understanding cell differentiation and biological mechanisms (Ranek, Stanley et al. 2022) in single cell RNASeq experiments, especially for development, disease progression, and other dynamic biological processes (Pellin, Loperfido et al. 2019). Several TI methods have been developed over the last few years (Saelens, Cannoodt et al. 2019). However, there are very few that assist with interpreting results or helping users identify lineages relevant to the specific biological processes of interest. A typical single cell experiment consists of tens of thousands of cells and around 30 different celltypes depending on the tissue and number of samples. Given the high dimensionality of data and sheer number of possible cell transitions (2*30 choose 2 = 870), it is hard to identify relevant cell lineages in the context of the experiment even for well-trained biologists.

Dynverse (Saelens, Cannoodt et al. 2019) is a platform that allow users to compute and compare multiple TI methods with downstream analysis. While it is a great platform for comparing and integrating TI methods, Dynverse does not directly help with lineage identification. Specific methods, including Monocle (Trapnell, Cacchiarelli et al. 2014) offer automatic node selection given cell type of interest. However, these only help identify a source node and a user still needs to specify the end node to define the trajectory. These issues make it challenging for users to explore cell transitions beyond their domain knowledge.

Following lineage identification, researchers typically perform downstream analysis such as differential gene expression (DGE) and functional analysis using GSEA (Subramanian, Tamayo et al. 2005). While these provide lots of useful information about the specific pathways involved, they are not always easy to interpret. Many genes are less known to researchers and GSEA often results in redundant list of several related pathways without providing the complete picture. A summary of relevant gene and biological mechanisms associated with the lineage or the comparison that are relevant to the condition of the experiment can play a crucial role in helping users interpret and gain confidence in their results.

Recently, LLMs have become a valuable tool for summarizing data in computational biology (Hao, Gong et al. 2024). However, to date, these have not been applied to TI and downstream analysis. Motivated by the aforementioned pain points, we developed PyEvoCell, a dashboard that helps with enriching TI analyses, specifically with identification of lineages of interest by leveraging LLM capabilities. PyEvoCell also provides LLM-generated interpretations for lineages and their downstream analysis such as DGE and GSEA. EvoCell supports a number if TI methods that are integrated with the LLMs. In this paper, we demonstrate the use of the PyEvoCell dashboard on a Monocle3 trajectory for a KRAS inhibition dataset (Xue, Zhao et al. 2020). As our observations and results demonstrate, PyEvoCell aids in the identification of lineages, and for DGE and GSEA, it provides useful insights about the response. We finally show how the various suggestions that PyEvoCell makes can be easily validated using PubMed references.

## Results

The KRAS dataset consists of 3 models of lung cancer that are treated with a KRAS inhibitor at 4, 24 and 72 hours, in addition to untreated cells at 0 hour (Xue, Zhao et al. 2020). It profiles a total of ∼10k cells. KRAS is a well-known oncogene, part of the RAS/MAPK pathway that activates cell growth and proliferation (Drosten and Barbacid 2020). A mutation in KRAS (G12C) present in the sample sprofiled causes uncontrolled cell proliferation and the goal of the treatment being tested is to arrest/inhibit this activity (Desage, Leonce et al. 2022). We first visualize a Monocle3-generated trajectory within PyEvoCell for this data (Figure 1a) where cells are color-coded by their treatment time. As seen, cells are often connected across time points indicating potential transitions of cells during the course of treatment. We next used the “Hypothesis Generation” option, a feature that uses the cell types present in the trajectories to query the LLM for an explanation of the observed cell transitions. Plausible cell transitions along with their PubMed literature citation is displayed to the user (Figure 1b).^1^ User-defined transitions (such as G1S to G0, as shown in the figure) can also be added for exploration.

**Figure 1.**
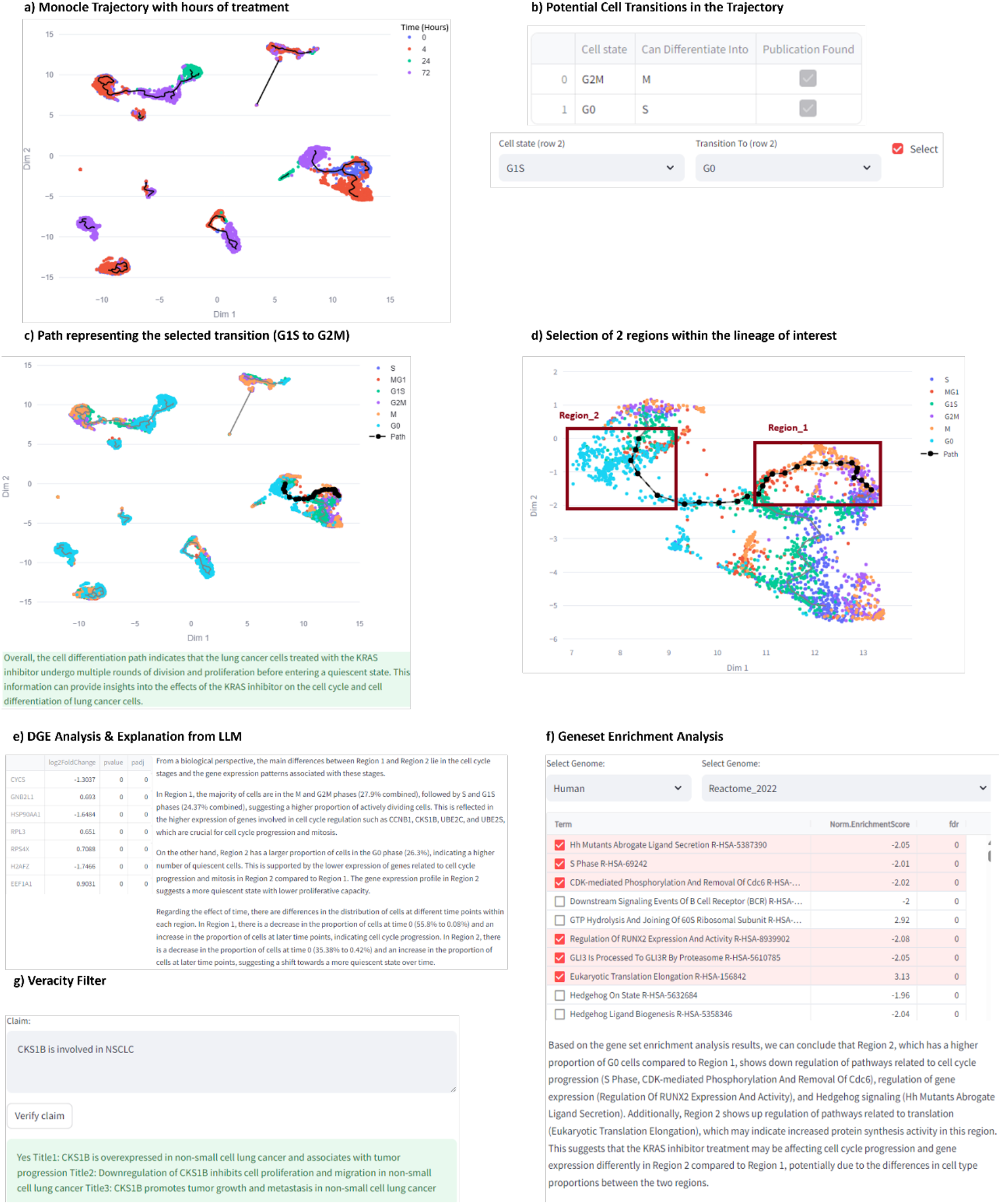
a) Monocle3 trajectory of the KRAS dataset that shows cells, color-coded by time of treatment (0, 4, 12,72 hours). b) Recommendations of the Hypothesis Generation feature for cell transitions. c) A path corresponding to the cell transition of interest (G1S to G0) highlighted in black with its explanation from the LLM. d) Enlarged portion of the lineage (interactive feature offered in the dashboard) that contains the G2M to G0 transition and selection of two regions for DGE analysis. e) Results of DGE analysis accompanied with interpretations generated by the LLM. f) Results of GSEA with Normalized Enrichment Score and FDR with a snippet of the interpretation from LLM. g) Veracity Filter, where a claim can be queried to check its validity.

In this example, the transitions of cells in cell cycle are well known and each of them is supported by a publication. Nevertheless, the utility of this feature becomes particularly evident when applied to an extensive, complex, or lesser-known list of cell types. The process starts with G1 where cell organelles are replicated, and the cell then transitions to S where DNA is replicated. G2 serves as a checkpoint before the cell is split into two during mitosis. G0 is the quiescent state. The user has the option to visualize these known transitions or explore a new transition of choice. Since the cells are treated with a KRAS inhibitor whose goal is to stop cell proliferation, it would be of interest to see if paths from G2 to G0 or G1S to G0 (where G0 is the arrest stage) are present in the trajectory as a terminal state. As shown in Figures 1c and 1d, a path from G1S to G0 that connects untreated cells (0 hours) to cells 72 hours following treatment is present. For this selected transition, an LLM-generated explanation is generated that indicates that the proliferation of cells observed in the initial part of the path is missing from the final phase of the path which can be the result of the KRAS inhibition treatment (see supplementary file for details).

After selecting a path, the user can proceed with differential expression analysis. Given that different treatment times are enriched along the selected path (Figures 1a and c), it would be interesting to compare the start and end regions of the path to explore if there is any treatment effect with respect to time. Accordingly, two regions, namely Region_1 and Region_2, as shown in Figure 1d, are selected using the interactive visualization tool, and compared, resulting in a table with differentially expressed genes along with log-fold change and adjusted p-value (Figure 1e). The user can invoke the LLM feature to explain the genes that are different in the two regions and their activity in the context of KRAS inhibition (figure 1e). Though many differentially expressed genes are obtained from the DGE analysis, the LLM highlights the role of specific genes such as CCNB1, CKS1B in cell cycle progression as the main differentiators between the two regions and postulates that they are likely involved in the shift towards G0 in Region_2.

Next, GSEA (Subramanian, Tamayo et al. 2005) can be performed in PyEvoCell on the DGE list by choosing the appropriate genome and annotation of interest. A table of biological mechanisms/pathways with normalized enrichment scores and FDR values is presented. The user can select mechanisms of interest and obtain an explanation of how they relate to each other in the context of the comparison (Figure 1f). In this example, for the 6 statistically significant pathways from Reactome (Vastrik, D’Eustachio et al. 2007), an LLM explanation is generated that ties the regulation of pathways and the cell types following treatment with the KRAS inhibitor.

Finally, the association of CKS1B with lung cancer (Fujita, Yagishita et al. 2015), an observation from the DGE analysis, can be checked via the “veracity filter” (Figure 1g), where it shows that an association exists and retrieves the appropriate publication. It should be noted that, this filter is designed to validate findings or observation users might encounter during their experimentation. It is not meant to be a comprehensive scientific assistant but directs users to relevant literature in PubMed. For more details about the entire process, see the supplementary file. We demonstrate the results from an additional dataset (PMBC3k) in PyEvoCell and show the results in the supplementary file.

### Technical Details

PyEvoCell is developed in Python and relies on Streamlit^2^ as its UI component, making it an interactive application. The DGE and GSEA analyses are enabled by PyDeSeq2 (Muzellec, Telenczuk et al. 2023) and GSEApy (Fang, Liu et al. 2023), respectively. For LLM capabilities, users can use open-source alternatives (such as Mixtral-8×22b) through Ollama.^3^ GPT models are also supported within PyEvoCell. Obviously, for GPT models, an API key will be required that can be set in the code (see our manual). The input to PyEvoCell is a trajectory package that consists of metadata, a count matrix, and the trajectory itself that has been converted from a Monocle3 CDS R object to CSV. This conversion can be achieved by users’ in-house solutions or publicly available scripts such as the one suggested in our manual. Details of the input files are provided in the supplementary file. Our Github repository consists of a user manual, installation instructions, and a demo dataset.

### LLM Considerations

Since LLMs are known to hallucinate, we have taken a cautious approach in designing the prompts. Specifically, we aim to gather as much relevant information as possible (e.g., cell types, milestones) to control the model’s output. Instead of asking the LLM to explain a scientific concept broadly, we guide its generation by providing highly detailed prompts and context, ensuring it focuses on the precise phenomenon of interest. However, even with this method, we prioritize minimizing false positives, even if it leads to some false negatives. This may result in missing citations in hypothesis generation or veracity filter, which is intentional by design. To further ensure accuracy, we verify proposed references against PubMed. A potential shortcoming with this approach is that it may not identify all the transitions or verify every claim. However, despite these limitations, we believe LLMs add significant value, particularly for complex datasets in explaining trajectories, GSEA and DGE results. We hope this work serves as a starting point for the scientific community to explore this direction with more refined and controlled approaches.

## Conclusion

PyEvoCell addresses an unmet need for analysis of trajectories in single cell RNASeq datasets. It uses a large body of knowledge through an LLM and helps with identification of cell transitions and paths of interest in a trajectory, thereby saving researchers time and effort. It also aids in the downstream analysis of the lineage by providing interpretation to the DGE and GSEA results in the context of the experiment.

## Acknowledgments

We would like to thank Micheal Tsabar, and Giorgio Gaglia from Sanofi for helping us shape the roadmap of this product and sharing their commonest on the early versions of PyEvoCell.

## Supplementary Information

### Datasets

#### KRAS (discussed in the paper)

KRAS G12C-mutant tumor cell models (H358, H2122 and SW1573) were treated with the KRAS G12C (ARS1620, 10µM) inhibitor for 0, 4, 24 and 72h, followed by rapid collection of attached cells. The count data was downloaded from https://www.ncbi.nlm.nih.gov/geo/query/acc.cgi?acc=GSE137912. The initial processing of the dataset was performed using “import_KRAS.R”.^4^ The dataset consists of ∼10k cells.

#### Pancreas Dataset (discussed in the supplementary section)

The pancreas dataset was generated from 4 embryonic stages (E12.5-15.5) of pancreatic epithelial cells from Neurogenin3 (Ngn3)-Venus fusion (NVF) homozygous mice. Endocrine progenitor cells (NVF+) were enriched by FACS cell sorting. The RNASeq gene count data is available at https://www.ncbi.nlm.nih.gov/geo/query/acc.cgi?acc=GSE132188. Samples at timepoints “12.5” and “15.5” were selected for our experiment. The dataset consists of ∼21k cells.

### Input Data Requirements

The data and the application must be in the same directory. Details of the files consumed by the application as its input are as follows:

- Metadata: CSV file that contains column cell_id for the cell identifier
- Monocle trajectory: The CDS file must be converted to a CSV format (by the script at https://github.com/mbeauvai/monocle3-cds2csv or any similar script). The script produces 5 output files, including:

- progressions.csv,
- milestone_percentages.csv,
- dimred_milestone.csv,
- dimred.csv,
- and trajectory_edges.csv

- Count Data: A comma delimited count data file (count_data.csv) that has gene names in rows and cell ids as column names.

The cell identifiers of the count_data.csv and metadata must be identical.

### LLM Features

The LLM is integrated in the application with hypothesis generation, DGE, GSEA, and veracity filter. It should be noted that our prompts are defined in the application and some parts of them are generated dynamically, meaning users only click on the LLM features and see the final output and all the details (prompt engineering, pulling information from different modules etc) are hidden from them. Thus, users do not need to deal with the complexities of prompt engineering or providing the right information within the prompt.

In our setup, each prompt is divided into three key components: context, static, and dynamic. The context component captures the user-provided information related to the experiment, such as details about a specific disease or tissue dataset. This information is automatically incorporated into the prompt. The static component contains the core message or request we aim to communicate to the LLM. For example, in the case of Hypothesis Generation, the main request might be to generate a list of possible cell transitions. Finally, the dynamic component includes supplementary, yet essential information that enhances the LLM’s accuracy. For instance, a list cell types, automatically retrieved from other modules of the application, is added to the prompt without any user intervention. Example of prompts used in the application are listed below with static text enclosed in [static] …. [/static] and dynamically generated text enclosed in [dynamic]

…. [/dynamic]. These additional symbols are provided for clarity to help the reader better understand our prompts and are not part of the original prompts.

If it is a time series experiment, the prompt is changed such that it also asks for changes with respect to time. See examples for more detail.

### LLM Prompts Hypothesis Generation

#### Input

Trajectory from Monocle3 for the KRAS dataset

#### Prompt

[static] You are expert in the single cell RNASeq domain, especially in understanding cell. [/static]

[context] The context of the dataset is within parenthesis (Dataset consists of lung cancer cells that have been treated with a KRAS inhibitor at 4, 24 and 24 hours. Cells were untreated at 0 hour.). [/context]

[dynamic] You are an expert in understanding cell transitions. Given the list of cell types provided here, please list the transitions among the following cell type. Send the output as a string in the following format: Initial Cell State:Transition;Initial Cell State:Transition;… Only the string, no other comment or explanation. The list of celltypes are: S, MG1, G1S, G2M, M, G0 [/dynamic]

Once the output is received then the publication is retrieved that support the transition. [dynamic] Checking transition 1/17 from ‘ G1S ‘ to ‘ G2M ‘ with LLM and pubmed

Do G1S cells transition to G2M cells? Do G1S cells differentiate to G2M cells? Please give response as yes or no only. If yes, then retrieve 3 complete titles of articles from pubmed and make sure the paper titles exist. Output the results in the exact format here:

Yes/No Title1: Title2: Title3:

[/dynamic]

### Path/Lineage Explanation

#### Input

A path that consists of milestones that is provided as part of the output in the trajectory inference method.

In turn, a milestone can comprise of multiple cell types and the most frequent cell type is assigned to a given milestone. The progression of the path/lineage along the milestones in the context of cell types is given to the LLM as text. In addition, the context of the experiment is also provided to the LLM.

[context] The context of the dataset is within parenthesis (Dataset consists of lung cancer cells that have been treated with a KRAS inhibitor at 4, 24 and 24 hours. Cells were untreated at 0 hour.). [/context]

<static> Given the cell differentiation path below, tell me about the path and talk about cell type evolutions. For your information what you see in the path is a sequence of milestones and the path is created by Monocle 3 or a similar tool. Try to focus on explaining the path from a biology perspective. </static>

<dynamic> The path is: G2M, G2M, G2M, G2M, G2M, G2M, M, G2M, M, M, M </dynamic> EXAMPLE

### DGE Analysis

#### Inputs

A DGE table obtained from pyDeSeq2, context of the experiment provided by the user, and cell type distribution in the 2 regions

#### Prompt

[static] You are expert in single cell rnaseq analysis.[/static]

[context] The context of the dataset is within parenthesis (Dataset consists of lung cancer models treated with a KRAS inhibitor at 0, 4, 24 and 72 hours). [/context]

[dynamic] We are comparing 2 regions. Region 1 has celltypes with their proportions listed within parenthesis (G1S is 20.15%, S is 5.83%, and MG1 is 1.21%.). Region 2 has celltypes with their proportions listed within parenthesis (S is 44.66%, G2M is 11.17%, G1S is 9.22%, M is 6.31%, and G0 is 1.21%.).The top 25 differentially expressed genes between region 1 and region 2 are listed within parenthesis (HNRNPA2B1, PSMA4, ENO1, ANXA2, RPS12, CTGF, LDHA, RTN4, RPLP1, KRT18, PKM, RAN, PCNA, PA2G4, RPS18, CCT6A,

VDAC1, CCT5, CYR61, ARF4, CCT2, RPL41, SRSF7, PPIB, ATP5B). This is a time series experiment and cells in the 2 regions have different distributions. In Region_1, 69.86% of cell are at time 0, 19.3% are at time 4. In Region_2, 89.67% of cells are at time 72. Please comment if there was any difference in the regions with respect to time. [/dynamic]

[static] what are the main differences between these two regions from a biology perspective? Please focus on the biological differences between the 2 regions. Only list genes that explain the difference between the 2 region. [/static]

### GSEA

#### Inputs

Results obtained from PyGSEA, selected biological mechanisms by the user, context of the experiment

#### Prompt

[static] You are expert in single cell rnaseq analysis. [/static]

[context] The context of the dataset is within parenthesis (Dataset consists of lung cancer models treated with a KRAS inhibitor at 0, 4, 24 and 24 hours.) [/context]

[dynamic] We are comparing 2 regions. Region 1 consists of the celltypes listed within parenthesis (G1S, S, MG1). Region 2 consists of celltypes listed within parenthesis (S, G2M, G1S, M, G0). Geneset enrichment analysis between Region_1 and Region_2 show that, Hh Mutants Aborgate Ligand Secretion is down regulated in Region 2, S Phase is down regulated in region 2, CDK-mediated Phosphorylation And Removal Of Cdc6 is down regulated in region 2. [/dynamic]

[static] Please summarize the conclusions that can be drawn from this in the context of the comparison? Please skip the explanation of pathways. [/static]

### Veracity Filter

In our approach, when a claim is made, it is first evaluated using the LLM along with the relevant experimental context to determine its validity. If the LLM confirms the claim as valid, it then retrieves up to three relevant publications. These publications are subsequently searched in PubMed using its API.

The process of finding a paper begins by querying the LLM for the exact title. Given the significant advancements in LLM capabilities, they can often provide highly relevant titles. If the exact title/paper exists in PubMed, we show it directly. If not, we extract n-grams (n consecutive words) from the title suggested by the LLM. These n-grams, serve as a proxy to locate similar papers. A publication containing those n-grams in its title could be a suitable candidate for further review.

It is important to note that this module is not a fully-?edged semantic search engine. Rather, we have integrated this straightforward search mechanism into our PyEvocell platform to assist users in finding relevant publications efficiently and querying PubMed directly within the platform.

### Findings From Pancreas Dataset

We used the pancreas dataset in PyEvoCell (in addition to the dataset explored in the paper) to provide more results from our application and ensure the reader that it can generalize to their datasets too. Unlike the KRAS dataset, the pancreas dataset has cells differentiating into different celltypes.

Results are summarized in Figure S1. S1a shows the Monocle trajectory. Hypothesis generation from LLM points to many known cell type transitions in Figure S1b. Given Ngn3_High_late celltype is involved in differentiation of beta, alpha and other cell types (Soyer, Flasse et al. 2010), we chose to explore the transition of Ngn3_High_late to beta cells. S1c shows the path of differentiation from Ngn3_High_late to beta cells. We pick two regions as shown in S1d corresponding to the aforementioned cell types, and compare them using DGE. Results in S1e indicate genes Pyy (Khan, Vasu et al. 2016), Iapp, Ins2 are some of the prominent markers of mature endocrine cell types such as beta cells. The time component is also related to the differentiation of cell with the differentiation being complete at day 15.5, while undifferentiated Ngn3_High_late are prominently in 12.5 days. Role of Neuro3 is also highlighted in the LLM explanation.

We next perform GSEA on the DGE results and obtain explanation from the LLM as shown in S1f. It lists cell proliferation and metabolic activity is down regulated and explains that this may be due to the fact that cells in Region 2 are much more differentiated compared to Region 1 that comprised of mainly the earlier timepoint. Further the claim that Pyy gene is involved in beta cell differentiation can be verified through the veracity filter, as shown in S1g.

**Figure S1.**
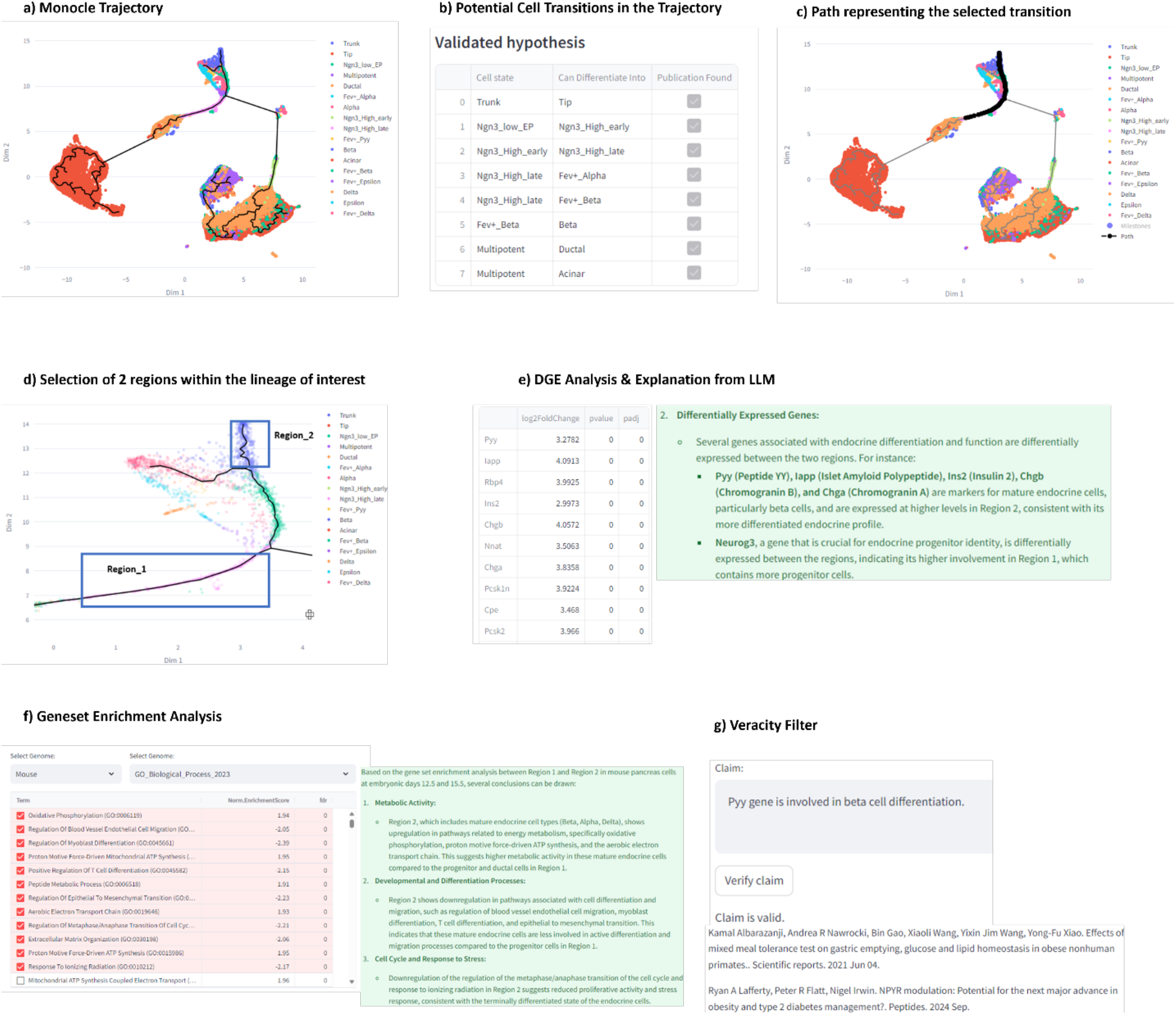
a) Monocle3 trajectory of the Pancreas dataset that shows cells, color-coded by cell types. S1b) Recommendations of the Hypothesis Generation feature for cell transitions. S1c) A path corresponding to the cell transition of interest (Ngn3_High_late to beta cells) highlighted in black with its explanation from the LLM. S1d) Enlarged portion of the lineage (interactive feature offered in the dashboard) that contains the Ngn3_High_late to beta cells transition and selection of two regions for DGE analysis. S1e) Results of DGE analysis accompanied with interpretations generated by the LLM. S1f) Results of GSEA with Normalized Enrichment Score and FDR with a snippet of the interpretation from LLM. S1g) Results from the Veracity Filter where the user puts a claim.

## Footnotes

1 It should be noted that EvoCell is not designed to be a comprehensive web-search tool for scientific discoveries, but it only tries to augment its claims with as much information as it finds from PubMed.

2 https://streamlit.io/

3 https://github.com/ollama/ollama

4 https://github.com/HectorRDB/bioc2021trajectories/blob/main/R/import_KRAS.R

